# Genetic tools that target mechanoreceptors produce reliable labeling of bladder afferents

**DOI:** 10.1101/2022.09.03.506472

**Authors:** Emily L. Tran, Sara A. Stuedemann, Olivia D. Link, LaTasha K. Crawford

**Affiliations:** Department of Pathobiological Sciences, University of Wisconsin-Madison School of Veterinary Medicine

**Keywords:** urinary bladder, mechanoreceptors, afferents, sensory neurons, whole-mount immunohistochemistry

## Abstract

Mechanosensitive bladder sensory neurons are critical for sensing bladder distention, but their role in bladder pain and bladder pathology is poorly understood, due in part, to the challenges of identifying mechanoreceptors in tissue sections. A lot of what is known about how disease alters sensory innervation of the bladder comes from studies that traditionally focus on nociceptive nerve terminals. In seeking tools to characterize the role of non-nociceptive afferents in the bladder, we first examined neurofilament heavy (NFH), a marker for medium to large-diameter myelinated A fibers, in combination with the common marker for peptidergic nociceptors, calcitonin gene-related peptide (CGRP). While there was partial overlap between NFH and CGRP, 87% of NFH fibers were CGRP-negative, underscoring the abundance of non-nociceptive A fibers nerve terminals in the bladder. Two mouse lines that have been used for genetic labeling of mechanoreceptors of the skin were tested for their ability to label bladder afferents. Once crossed to Cre-dependent reporter lines, tyrosine kinase B (TrkB) TrkB^CreER2^ mice can be used to label A-delta mechanoreceptors while receptor tyrosine kinase ret protooncogene (Ret) Ret^CreER2^ mice can label a combination of A-beta mechanoreceptors and non-peptidergic nociceptors in adult mice (late Ret). Both mouse lines produced successful labeling of bladder nerve terminals demonstrating partial overlap with NFH and minimal overlap with CGRP. Thus, we have identified new genetic strategies to investigate CGRP-negative subpopulations of bladder afferents that remain largely uncharacterized in studies that target peptidergic nociceptors. These tools can help elucidate the role of mechanosensitive afferents in bladder pathophysiology and urologic chronic pain.

## Introduction

Bladder sensation is mediated by primary sensory neurons, also termed afferents, whose cells bodies are found in dorsal root ganglia (DRG) that sit bilaterally at each vertebral level next to the spinal cord. Additional sensory innervation is thought to originate from visceral afferents of the vagus nerve (Herrity et al., 2014; Jancsó & Maggi, 1987). A simplified view divides bladder afferents into mechanoreceptors, mechanosensitive afferents which respond to lower degrees of stretch or bladder “fullness”, and nociceptors, which respond to excessive distention, irritants, and noxious stimuli. It is thought that distinct populations of mechanosensitive afferent fibers sense stretch in the bladder wall and send signals to the central circuits, where relaxation of the bladder wall is then mediated by specific pathways to accommodate more fluid (Fowler et al., 2008; Sugaya et al., 2005). Once the pressure threshold is reached, the activity of these pathways is temporarily inhibited, allowing contraction of bladder wall and relaxation of external sphincter muscles, resulting in voiding (Fowler et al., 2008; Keast et al., 2015). Because mechanoreceptors can encode stretch and distension into the noxious range, it is theorized that under pathologic conditions, mechanoreceptors may contribute to the discomfort and pain experienced in bladder disease.

More detailed understanding of bladder afferents has been gained through the study of the electrophysiological properties of unmyelinated slowly conducting C fibers, largely considered nociceptors, and moderately myelinated, fast-conducting A-delta fibers that are largely assumed to be mechanoreceptors (Birder et al., 2010; Keast et al., 2015). It is theorized that A-delta fibers are typically responsible for firing at lower bladder pressures. As the bladder fills, the C fibers are recruited, causing the sense of urgency. As pressure increases, C fibers can cause the sensation of bladder pain (Grundy et al., 2019). However, in models of bladder inflammation, both C fibers and mechanosensitive fibers become hypersensitive suggesting that mechanoreceptors may also play a role in bladder pain (Xu & Gebhart, 2008; Zagorodnyuk et al., 2007). Reliance on electrical properties or response properties to identify mechanoreceptors is fraught with challenges, if these defining properties are altered by the disease process under study. Though studies often use measures of conduction velocity to define which subtypes are present, conduction velocity can overlap between some subtypes and somatosensory pain conditions can lead to high variability in conduction velocity or an increase in the number of neurons that cannot be categorized by conventional electrophysiology methods (Boada et al., 2015; Wang et al., 2016). This underscores a critical need for better tools to enable the study of mechanosensitive bladder afferents and their role in bladder pain and other diseases.

A major hurdle in the study of mechanoreceptors and their role in bladder disease is the historical lack of reliable immunohistochemistry (IHC) markers for mechanoreceptor subtypes. Nociceptors are a common focus of studies of bladder disease, where markers like calcitonin gene related peptide (CGRP) can be used to identify the peptidergic subtype of nociceptors. However, mechanoreceptors have been largely overlooked in molecular studies of bladder disease due to our inability to identify them in tissue sections. In this study, we attempted to identify mechanosensitive afferents in the bladder by first examining a non-specific marker of myelinated axons, as the vast majority of mechanoreceptors are moderately to heavily myelinated. After confirming a large population of CGRP-negative, myelinated neurons that innervate the bladder, we used genetic strategies known to label specific mechanoreceptor subtypes in the skin (reviewed in Crawford and Caterina 2020) to investigate patterns of bladder afferent innervation. Whole-mount fluorescent IHC identified distinct patterns of innervation with a notable lack of co-localization with marker of peptidergic nociceptors, highlighting new tools to interrogate bladder afferents that are largely uncharacterized in bladder disease.

## Methods

### Animals

All experiments were approved by the Institutional Animal Care and Use Committee of the University of Wisconsin Madison. Animals were housed in groups of 2 to 4 in a temperature-controlled room with and 12:12 hr light/dark cycle and ad libitum food and water. Genetic labeling of neuronal subpopulations was generated as previously described by crossing Rosa26LSL-tdTomato (strain Ai14) (Madisen et al., 2010) to TrkB ^CreER2^ (Rutlin et al., 2014), NtnG2 N’Cre Abhd3 C’Cre Split Cre intein BAC transgenic line (Split-CRE) mice (Fleming et al., 2016; Rutlin et al., 2014) or by crossing Rosa26LSL-tdTomato (strain Ai9) mice to Ret^CreERT2^ (Luo et al., 2009). TrkB ^CreER2^;Rosa26^LSL-tdTomato^, Ret^CreER2^;Rosa26^LSL-tdTomato^, and Split-CRE;Rosa26^LSL-tdTomato^ offspring are referred to hereafter as TrkB, Ret, and Split, respectively. Transgenic lines were maintained on a genetic background consisting predominantly of C57BL/6 with contributions from other strains prior to back-crossing efforts. The dams and sires used to generate offspring for this study expressed additional knock in alleles at the Tau or Advillin locus that were not utilized for these experiments. Cre expression was induced with administration of 2mg/kg of Tamoxifen in wheat germ oil by oral gavage to Ret line offspring at 3-6 weeks of age (adult/late Ret) and to TrkB line offspring at 3-5 weeks of age. Both male and female mice were used at 3-8 months of age, including 4 Ret mice and 4 TrkB mice. Three Split mice at 3-4 months of age were also examined.

### Tissue Collection and Processing

Animals were deeply euthanized with a lethal dose of intraperitoneal Euthasol. Bladders were dissected, a ∼1mm slit was cut into the dome, and the bladder was emptied completely before overnight immersion fixation in 4% paraformaldehyde at 4°C. Two TrkB bladders were collected after perfusion fixation but previous analysis determined that they had identical labeling patterns to those that were immersion fixed (data not shown).

### Whole-mount Immunohistochemistry (IHC)

Whole-mount staining was adapted from a published protocol for skin (Li & Ginty, 2014). Briefly, bladders underwent a series (10-14) of 20 min washes in a solution of 1% Triton-X in PBS before and after a 48-hour primary antibody incubation at room temperature. A separate, overnight room temperature incubation in secondary antibody was followed by ten 30 minute washes in 1% TritonX in PBS, then 5 washes in PBS. Multiplex antibody cocktails were made in a whole-mount blocking solution consisting of 5% heat activated normal donkey serum (Sigma Millipore), 20% Dimethyl sulfoxide (Sigma Millipore), and 1% Triton-X in PBS. The antibodies used are summarized in Table 1. An antibody recognizing mCherry was used to amplify the transgene-mediated tdTomato labeling. After incubation, bladders were opened with a longitudinal midline cut, splayed open, mounted lumen-side down, and coverslipped with Fluoromount G mounting media (refractive index = 1.4). Previous experiments showed comparable results from tissues processed with this technique and tissues dehydrated and chemically cleared using one part Benzyl Alcohol and two parts Benzyl Benzoate (BABB) (Supplemental figure 1). This indicated that BABB clearing of our samples was unnecessary given the deconvolution capabilities of the computational clearing algorithms used in this study (see Fluorescent imaging below). Thus, the bladders for this study were not chemically cleared prior to imaging.

**Table 1.** Antibodies used in this study.

| Antibody | Manufacturer | Catalog Number | RRID | Conc. |
| --- | --- | --- | --- | --- |
| Sheep Anti-CGRP Polyclonal Antibody, Unconjugated | Abcam | ab22560 | AB_725809 | 1:1000 |
| Chicken Anti-NFH Polyclonal IgY (200 kDa) Antibody | Aves Lab | NFH | AB_2313552 | 1:500 |
| Rat Anti-mCherry Monoclonal Antibody (16D7) | ThermoFisher | M11217 | AB_2536611 | 1:400 |
| Alexa Fluor 488-AffiniPure Donkey Anti-Sheep IgG (H+L) Antibody | Jackson ImmunoResearch Labs | 713-545-147 | AB_2340744 | 1:250 |
| Cy3-AffiniPure Donkey Anti-Rat IgG (H+L) Polyclonal Antibody | Jackson ImmunoResearch Labs | 712-165-153 | AB_2340666 | 1:250 |
| Alexa Fluor 647-AffiniPure Donkey Anti-Chicken IgY (IgG) (H+L) Antibody | Jackson ImmunoResearch Labs | 703-605-155 | AB_2340379 | 1:250 |

### Fluorescent Image Acquisition

Images of mounted bladders were acquired with an upright Thunder 3D Tissue Imaging Microscope DM6B-Z with motorized stage and 40 × 0.95 NA dry objective (Leica Microsystems) along with LAS X software (Leica Microsystems). A standardized technique targeted the same regions of the dome, body, and neck across all samples (Supplemental Figure S1). Z-stacks were acquired by targeting the subepithelium and detrusor muscle layer, avoiding the urothelium during image acquisition. The average thickness of the Z-stack was 90um including an average of 138 optical slices sampled at Nyquist frequency (0.293μm step size). LED light intensity and exposure times were optimized for each channel and image parameters were kept consistent across all samples.

Prior to analysis, pseudocolors were applied to each channel along with consistent, linear adjustments to the histogram. All images were deconvoluted and computationally cleared using the Thunder Large Volume Computational Clearing algorithm in LAS X. Thunder Good’s Roughness regularization for channel 1 (NFH) was 4.26e-6 and strength was 9.8e-1. Both channel 2 (CGRP) and channel 3 (mCherry) regularization was 4.26e-9 and strength was 9.2e-9 (Figure S1). To produce figures, images represent maximum projections of the acquired Z-stacks with a uniform 20% increase in contrast and brightness.

### Image Analysis

All quantitative analysis was done using NIS Elements-AR software (Nikon). Analyzed images included Z-stacks cropped to a consistent thickness of 80 Z slices, roughly corresponding to the detrusor muscle layer. From LAS X, .LIF files were converted to .TIFs and imported as .ND2 files into NIS Elements-AR. Measurements had to be calibrated from pixels (px) to micrometers which consisted of a 0.2μm to 1 px conversion. Rolling ball background subtractions set to 0.025um were applied to maximum projection images in each channelThresholding settings were determined for each channel and were consistent across all images. Binary area fraction of the total image area, (area of signal above threshold) / (total area analyzed), was quantified in maximum projection images as a measure of fiber density.

For colocalization analysis, rolling ball background subtraction was set to 0.004μm (50px) and applied to each channel for analysis of the entire Z-stack. Subtraction masks highlighting image objects 0.01μm (20px) or larger, were made to highlight large nerve bundles to remove them colocalization analysis. Regions of interest (ROIs) were produced using software-generated binaries that excluded objects within the subtraction mask. Colocalization then was determined using the intersection of binary masks created using consistent threshold settings optimized for each channel; binary areas within ROI and the ROI area were the key measurements acquired from each image in each Z-stack. Binary area fractions for colocalization were calculated as (binary area) / (ROI area x 80) where 80 is the total number of Z slices analyzed in each stack. Colocalized pixels of marker A + B, as a percent of total pixels of marker A were calculated from raw data using Prism (version 8.4.0, GraphPad).

### Statistical Analysis

Values are reported as the average ± the standard error of the mean (SEM) unless otherwise noted. All statistics were performed in Prism. Statistical comparisons were made using two-tailed unpaired t-tests and differences were assessed at a significance level of p < 0.05.

## Results

### NFH partially co-localizes with CGRP, but primarily labels CGRP-negative fibers

Because the vast majority of mechanoreceptors are generally characterized by moderate to heavy myelination, we first examined a non-specific marker of myelinated sensory axons to determine if there were CGRP-negative, myelinated nerve terminals in the bladder. Whole mount multiplex IHC of the mouse urinary bladder was used to evaluate axon terminals labeled with the 200kDa, heavy isoform of neurofilament (NFH) a marker of moderate to heavily-myelinated axons in the mouse (Dhandapani et al., 2018; Foster et al., 1967; Le Pichon & Chesler, 2014; Luo et al., 2009), that is thought to label A-delta fibers in the bladder (De Groat & Yoshimura, 2009; Lawson et al., 1993). We also stained for CGRP to label nociceptor fibers. Both NFH and CGRP were present in large nerve bundles and individual fibers throughout the dome, body, and neck of the bladder (Figure 1A). For both markers, variable branching patterns can be seen across locations, resembling what has been previously reported for CGRP-positive nerve terminals in the urinary bladder (Forrest et al., 2014; Sharma et al., 2020; Spencer et al., 2016). Quantitative analysis of innervation density of NFH-positive and CGRP-positive nerve terminals revealed that NFH-positive nerve terminals were in fact significantly more dense, (Figure E, p < 0.0001 using a two-way ANOVA, N = 8 mice). A closer look at bladder regions revealed NFH-positive nerve terminals to be more dense than CGRP in the dome (p = 0.002), body (p = 0.0014), and neck (p = 0.0279 using paired two-tailed t-tests, N = 8 mice). Surprisingly, our data reveals that some individual CGRP-positive axonal fibers co-expressed NFH (Figure 1C, i), as dual-labeled nerve terminals were found in all regions of the bladder. Colocalization analysis showed that of the CGRP-positive nerve terminals, 58.7% ± 5.5% co-express NFH, suggesting that a portion of these fibers are myelinated A-delta fibers rather than C fibers. We also found that 87 ± 4.1%, of NFH-positive nerve fibers were CGRP-negative (Figure 1C, ii). We reasoned that at least some of the NFH-positive, CGRP-negative nerve terminals in the bladder are likely to be mechanoreceptor afferents.

**Figure 1.**
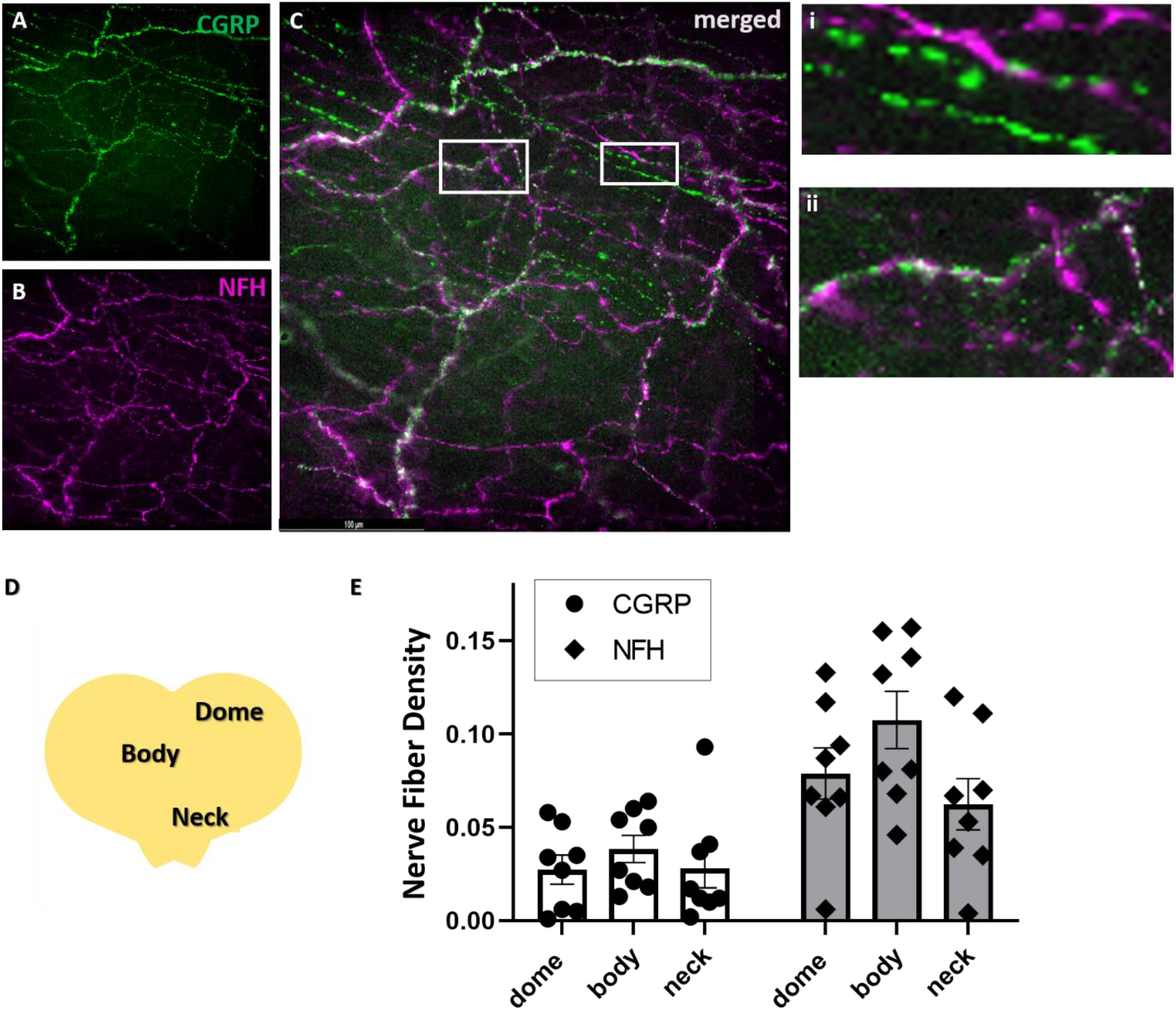
NFH-positive nerve terminals demonstrate partial colocalization with CGRP in the bladder. (A) CGRP-positive (green) nerve terminals are composed of large nerve bundles (not shown) along with small individual nerve fibers that occasionally have a beaded appearance. (B) NFH-positive (magenta) nerve terminals are composed of large nerve bundles (not shown) along with small individual nerve fibers (C) Merged image of CGRP and NFH, where many nerve fibers are singly-labeled (i) while other nerve fibers demonstrate colocalization (white), as show in an inset (ii). (D) Diagram showing the bladder regions targeted for quantitative analysis. (E) Quantification of nerve fiber density (binary area fraction) of CGRP and NFH-positive nerve fibers across the dome body and neck of the bladder show that NFH innervation is more dense that CGRP innervation (**p= 0.001, two-way ANOVA, N = 8 mice).

### Genetic labeling of bladder afferents using Ret and TrkB mouse lines

To circumvent the limitations of IHC, we used genetic strategies to try to identify mechanoreceptor afferents in the bladder. Transgenic mouse lines used for the study of mechanoreceptors in the skin were screened to determine which approach can produce genetically labeled neurons that innervate the urinary bladder. Ret expression in adult mice (late Ret) is characteristic of A-beta mechanoreceptors, C mechanoreceptors and nonpeptidergic nociceptors (Luo et al., 2007; Stucky & Lewin, 1999; Usoskin et al., 2014). TrkB expression in skin is typically specific to A-delta mechanoreceptors (Li et al., 2011; Rutlin et al., 2014). Genetic labeling of bladder afferents was successful following tamoxifen-induced Cre expression in TrkB ^CreER2^;Rosa26^LSL-tdTomato^ (TrkB) offspring and in Ret^CreER2^;Rosa26^LSL-tdTomato^ (Ret) offspring. The fiber density of labeled nerve terminals varied slightly across the region imaged in bladders from both mouse lines (Figure 2), with a slight increase in the body of the bladder for bot hTRk-B- and Ret-labeled afferents (Figure 2C). Ret-labeled nerve fibers of the neck were comparable to the dome and body while, TrkB-labeled nerve fibers in the neck were less dense than the body (Figure 2C; p=0.042, Friedman non-parametric ANOVA test, *p=0.040 body vs neck Dunn’s multiple comparisons test).

**Figure 2.**
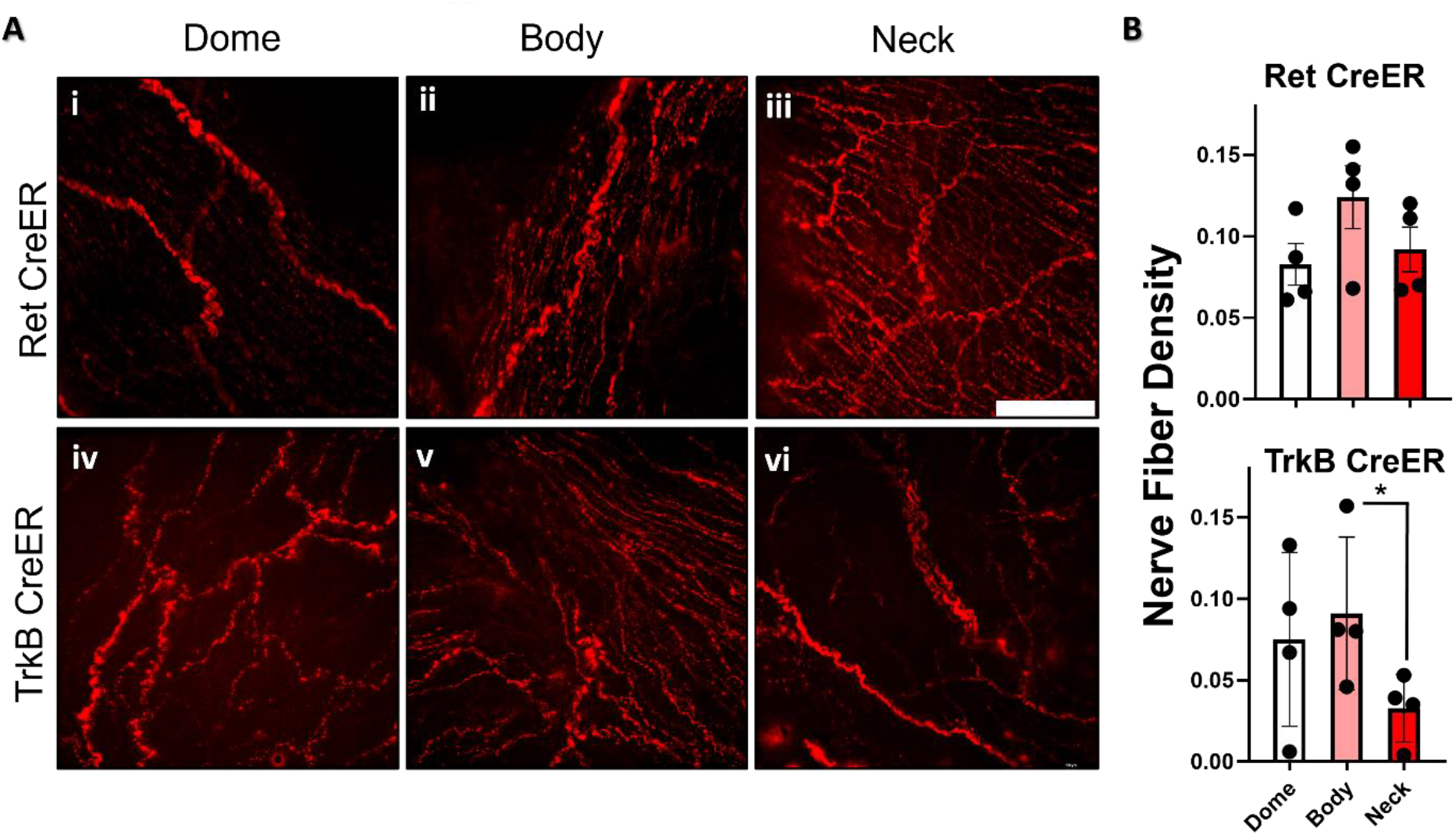
Cre-dependent tdTomato-labeling of nerve terminals is evident throughout the bladders of Ret and TrkB mice. (A) In Ret mice there are tdTomato-labeled nerve bundles and individual nerve fibers across the dome (i), body (ii), and neck (iii) of the bladder. (B) Likewise, in TrkB mice there are tdTomato-labeled nerve bundles and individual nerve fibers across the dome (i), body (ii), and neck (iii) of the bladder. (C) Quantification of fiber density (binary area fraction) including large nerve bundles and individual fibers show a slight increase in innervation density in the body of the bladder compared other regions. Ret-labeled nerve fibers of the neck were comparable to the dome and body while, TrkB-labeled nerve fibers in the neck were less dense than the body (p=0.042, Friedman non-parametric ANOVA test, *p=0.040 body vs neck Dunn’s multiple comparisons test). N = 4 per group. Whole-mount mCherry immunostaining for tdTomato-labeled axons. Scale bar is 100um.

### TrkB-labeled nerve terminals are CGRP-negative and consistent with A-delta mechanoreceptors

Further analysis sought to characterize the types of nerve terminals labeled by our genetic strategies. Multiplex whole-mount IHC of TrkB bladders were analyzed to determine the relative colocalization of labeled nerve terminals with CGRP and NFH. We limited our colocalization analysis to the body of the bladder to avoid the large nerve bundles that traverse the neck of the bladder enroute to other regions. We further excluded large nerve bundles from analysis by creating masks that systematically removed large bundles from the analyzed regions of interest, focusing primarily on individual nerve fibers for quantification of colocalization (Figure 3). Analysis revealed that 3.7% ± 1.4% of TrkB-positive afferents colocalized with CGRP (Figure 3C, bottom), indicating that the vast majority of TrkB-labeled nerve terminals are distinct from peptidergic sensory neurons. Approximately 37% ± 20% of TrkB-labeled fibers expressed NFH in the body of the bladder (Figure 4C, bottom), suggesting that many of these nerve terminals are myelinated, a characteristic feature of A-delta mechanoreceptors in other tissues.

**Figure 3.**
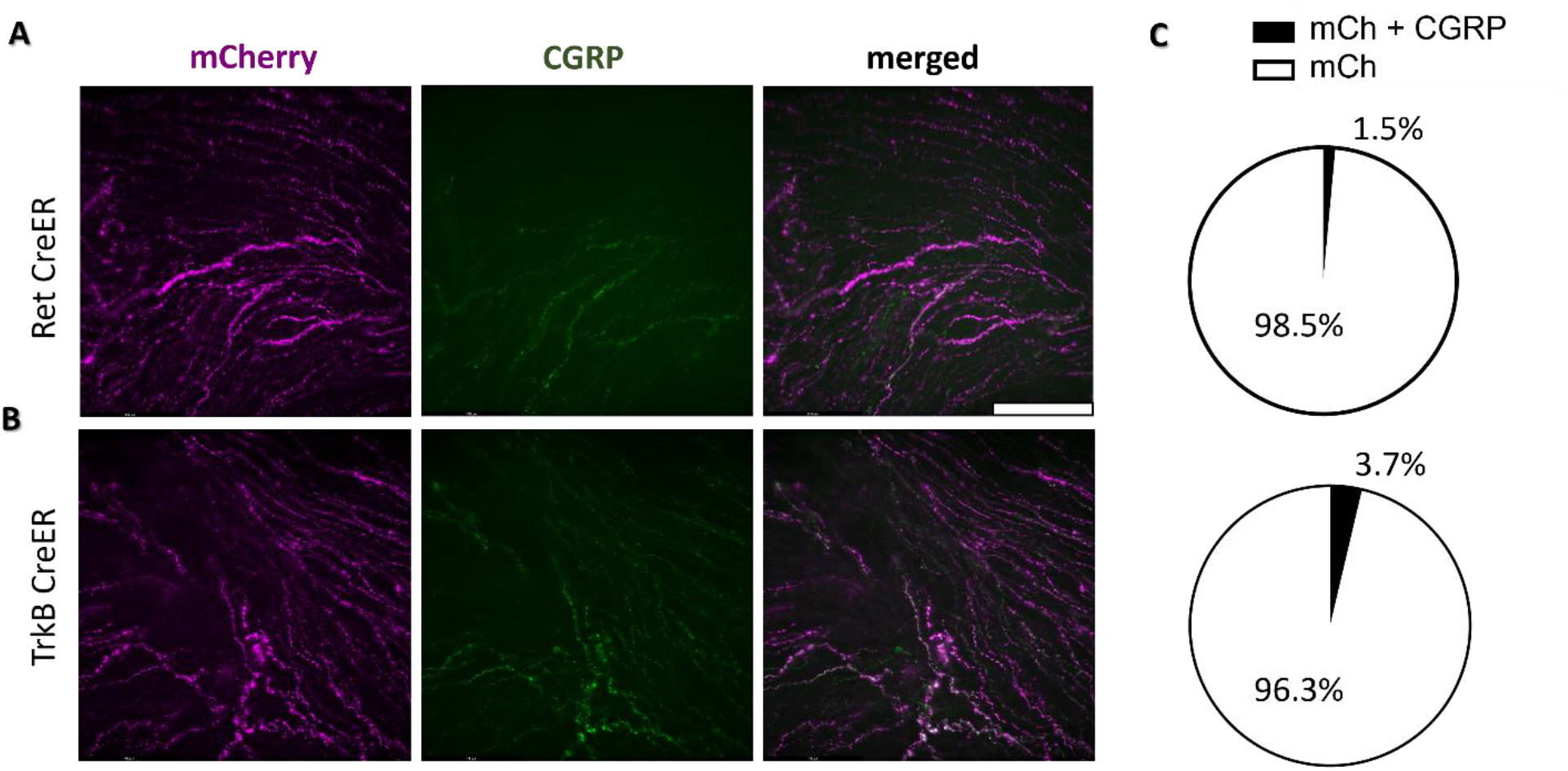
Genetically-labeled nerve terminals in the bladder of Ret and TrkB mice do not colocalize with CGRP. (A) In the bladder from a Ret mouse, tdTomato-labeled nerve terminals (magenta) do not colocalize with CGRP-positive nerve terminals (green). (B) In the bladder from a TrkB mouse, tdTomato-labeled nerve terminals are likewise largely independent of CGRP. Although images show maximum projections, quantification was performed on optical slices from Z-stacks. (C) Quantification of immunomarker colocalization in Ret bladders (N=4, top) shows that only 1.5 ± 1.0% of tdTomato-labeled fibers also express CGRP (N=4, top) while in TrkB bladders only 3.7 ± 1.4% of tdTomato-labeled fibers co-express CGRP. Whole-mount mCherry immunostaining for tdTomato-labeled axons, body region of the bladder. Scale bars are 100um.

**Figure 4.**
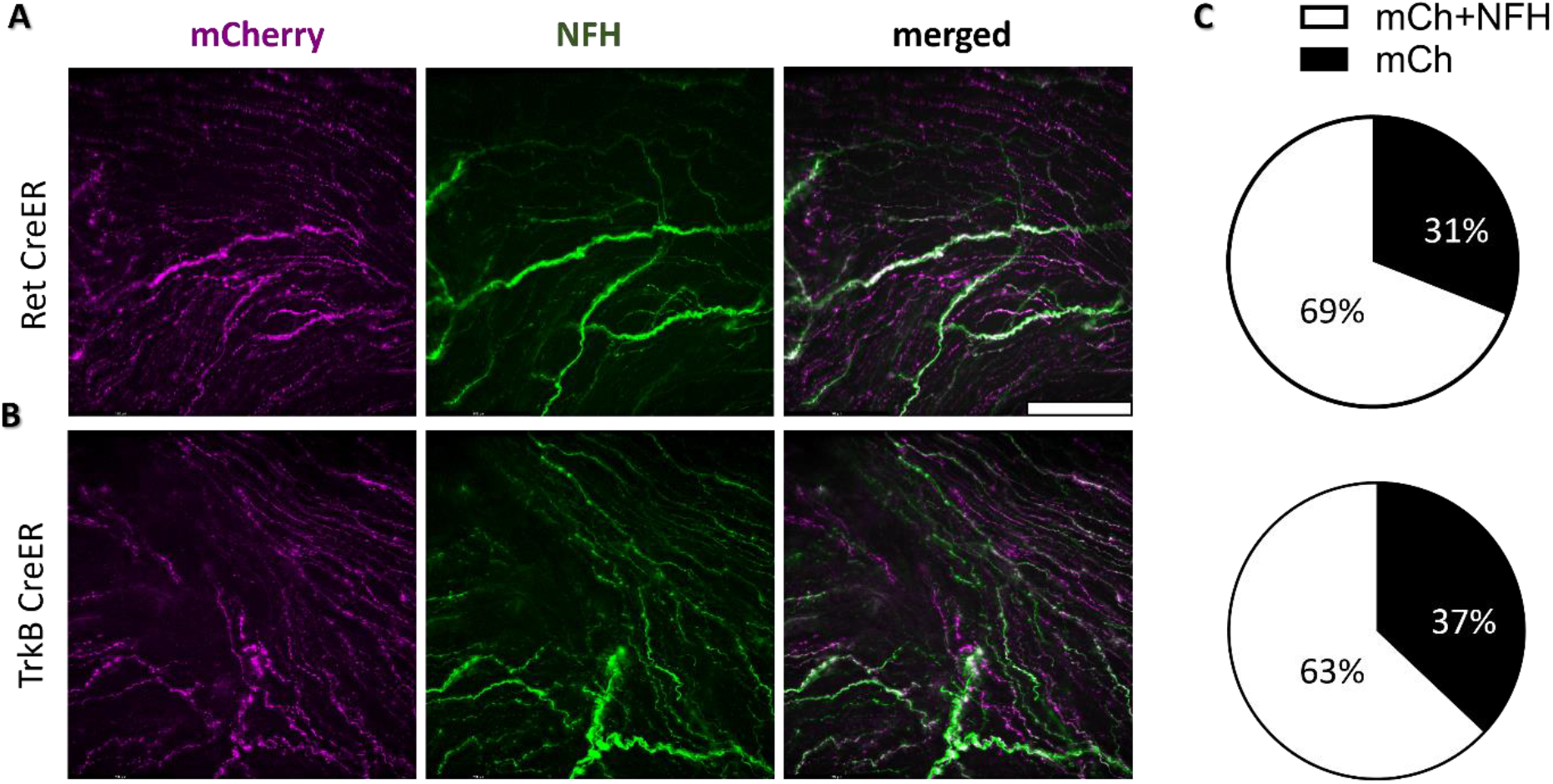
Genetically-labeled nerve terminals in the bladder of Ret and TrkB mice partially colocalize with NFH. (A) In the bladder from a Ret mouse, tdTomato-labeled nerve terminals (magenta) show partial overlap with CGRP (green). (B) In the bladder from a TrkB mouse, tdTomato-labeled nerve terminals likewise show partial colocalization with CGRP. Although images show maximum projections, quantification was performed on optical slices from Z-stacks. (C) Quantification colocalization in Ret bladders (n=4, top) shows that 31 ± 2.2% of tdTomato-labeled fibers also express NFH (N=4, top) while in TrkB bladders 37 ± 20% of tdTomato-labeled fibers co-express NFH. Whole-mount mCherry immunostaining for tdTomato-labeled axons, body region of the bladder. Scale bars are 100um.

### Ret-labeled nerve terminals partially colocalize with NFH

Quantitative colocalization analysis targeting individual nerve fibers revealed that Ret bladders express partial colocalization with NFH but minimal colocalization with CGRP. Of the total tdTomato-labeled in the body of the bladder of Ret mice, 1.5% ± 1.0% of nerve terminals also express CGRP (Figure 3A, 3C top). This was quite small compared to the 31% ± 2.2% of tdTomato-labeled fibers that are also NFH-positive (Figure 4A, 4C, top). Interestingly, of the few tdTomato-labeled, CGRP-positive fibers, 82% ± 4.8% also expressed NFH. This pattern of triple positive expression seen in 1-2% of the Ret-labeled fibers is consistent with peptidergic A-delta fibers. However, the overwhelming majority, 92% ± 4.6%, of tdTomato-labeled NFH-positive fibers are CGRP-negative, suggesting they are non-peptidergic myelinated A-fibers.

### Split-CRE labeled nerve terminals are minimal to inconsistent in the bladder

In the skin, Split-CRE induced tdTomato expression is seen in rapidly adapting A-beta low threshold mechanoreceptors (Fleming et al., 2016; Kuehn et al., 2019), though labeling patterns can vary as mice become aged (data not shown). For those mice whose skin innervation patterns reflect successful, specific labeling of rapidly adapting A-beta low-threshold mechanoreceptors (longitudinal lanceolate endings oriented around intermittent hair shafts), there was minimal to no innervation of the bladder (Supplemental Figure S2). This is consistent with the conventional understanding that A-beta mechanoreceptors do not innervate the bladder. However, one Split mouse demonstrated robust, aberrant, non-specific tdTomato labeling in the skin that was not specific to A-beta low-threshold mechanoreceptors. This mouse did show modest numbers of tdTomato-positive nerve terminals (Supplemental Figure S2) along with labeling of abundant unidentified non-neuronal cell populations in the bladder. This labeling was considered non-specific.

## Discussion

In order to advance the study of mechanosensitive innervation of the bladder, this study unearthed new strategies to identify non-peptidergic afferents in the mouse bladder. Using NFH as a marker of myelinated A-fiber types (De Groat & Yoshimura, 2009; Keast et al., 2015), we identified a large population of non-peptidergic A-fibers that comprised potential mechanoreceptors. We then identified two genetic strategies to identify bladder afferents, including TrkB-mediated expression of a fluorescent reporter and Ret-mediated expression of a fluorescent reporter, with conditional expression of the Cre recombinase induced at 3-5 weeks of age. The vast majority of afferents labeled using these two genetic strategies were CGRP-negative, confirming that they are non-peptidergic. Thus, despite the knowledge gained from a wide range studies of peptidergic nociceptive bladder innervation, the neuronal populations labeled by our genetic strategies remain largely uncharacterized by CGRP studies. Our data also demonstrated that the slight majority (58%) of CGRP-positive nerve fibers in the bladder were positive for the NFH marker of myelinated axons, suggesting that CGRP bladder afferents are not only C fibers, but also include A-delta fibers as well. This is consistent with previous reports of large diameter CGRP-positive neurons with retrograde tracers (Hwang et al., 2005) Interpretations that assume that CGRP innervation is synonymous and interchangeable with C-fiber sensory innervation may represent an inaccurate oversimplification of the heterogeneity of bladder innervation.

Inducible TrkB^CreER^ crossed to a Cre-dependent fluorescent reporter produced labeling of bladder afferents within each region of the bladder. At the timepoint utilized in our genetic strategy (∼3 weeks of age) TrkB expression in the skin is specific to A-delta mechanoreceptors (Li et al., 2011; Rutlin et al., 2014) whose nerve terminals form longitudinal lanceolate endings around a subset of hair follicles in haired skin (Abraira & Ginty, 2013; Crawford & Caterina, 2020; Li et al., 2011). A portion (∼37%) of TrkB-labeled nerve fibers in the bladder expressed the A-fiber marker NFH and were therefore consistent with A-delta mechanoreceptors. Further studies are needed to elucidate the identity of NFH-negative TrkB-labeled neve terminals.

Cre expression induced in adult offspring from Ret^CreER^ crossed to a Cre-dependent fluorescent reporter likewise produced labeling of bladder afferents within each region of the bladder. While early Ret is largely restricted to rapidly adapting A-beta mechanoreceptors, at the time point used in this study (tamoxifen administered to young adult mice), Ret expression can include rapidly adapting A-beta mechanoreceptors, A-beta field mechanoreceptors, non-peptidergic C nociceptors, and C low threshold mechanoreceptors (Luo et al., 2007; Stucky & Lewin, 1999; Usoskin et al., 2014). Our results also showed a proportion (31%) of Ret-labeled axons in the bladder expressed NFH. While this population of Ret-labeled neurons could be A-beta mechanoreceptors, the prevailing understanding of bladder innervation suggests that A-beta neurons do not innervate the bladder. This was supported by our findings that Split-CRE-labeled nerve fibers were essentially absent from the bladder in mice with confirmed, specific labeling of skin rapidly adapting A-beta mechanoreceptors (Fleming et al., 2016; Kuehn et al., 2019; Rutlin et al., 2014). Ret expression is required for the development and axonal migration of sympathetic neurons, but is downregulated in at least some autonomic subpopulations later in development (Enomoto et al., 2001; Ernsberger et al., 2020). However, less is known about how Ret expression changes over the life of the mouse in autonomic neurons that innervate the bladder. If Ret is expressed in adult autonomic motoneurons at 3-6 weeks of age (the time of tamoxifen administration in our study), it is possible that a subset of Ret-labeled NFH-positive, CGRP-negative nerve fibers in our study represent autonomic innervation, as NFH has been reported in neuronal cell bodies of the pelvic ganglia in the mouse (Forrest et al., 2014; Keast et al., 2015). The majority of Ret-labeled afferents were NFH-negative, likely include C-mechanoceptors or nonpeptidergic C nociceptors. C low-threshold mechanoreceptors are TH-positive, NFH-negative, unmyelinated sensory nerve fibers that originate from the DRG and can express Ret in the adult mouse (Li et al., 2011a; Usoskin et al., 2014). There are also TH-positive sensory neurons that originate in the DRG and innervate visceral organs including the urinary bladder (Brumovsky et al., 2012). Thus, it is quite likely that some of the NFH-negative, CGRP-negative Ret-labeled bladder afferents in our study were C-fiber mechanoreceptors. It is unclear whether Ret-driven labeling of bladder afferents can include labeling of non-peptidergic C-fiber nociceptors, as it does in the skin. The advent of strategies like single-cell sequencing and spatial transcriptomics throughout the developmental life span of the animal will be invaluable for elucidating molecular signatures of bladder-innervating sensory neurons. These data will help investigators hone the specificity of the Ret approach by directing induction of Cre to a particular time in development that achieves more specific expression patterns or by using immunomarkers whose colocalization with Ret-labeling can refine the targeted subpopulation.

The use of whole-mount staining techniques allowed us to examine and quantify intact nerve terminals as they traverse different layers of the bladder. Image acquisition using automated Z-stage microscopy paired with advanced deconvolution software permitted visualization of fine nerve fibers throughout the entire thickness of the tissue, at a resolution beyond what can be visualized by routine microscopy in the face of significant light scatter (Supplemental Figure S1). This permitted quantification of nerve terminal distribution patterns across three bladder regions in whole mount bladders, rather than qualitative or semi-quantitative descriptions of individual cross sections as has been conducted in other studies (Gabella, 2019; Rahnama’i et al., 2017). In Z-stack images collected from the suburothelium and muscle layers, we found a comparable to higher innervation in the body of the bladder compared to the bladder neck, across several subtypes of nerve terminals. This was distinct from other studies where the highest concentration of CGRP afferent nerve fibers has been reported in the bladder neck, particularly within the suburothelium (Gabella, 2019; Rahnama’i et al., 2017). The pattern of innervation by TrkB-labeled afferents and Ret-labeled afferents may differ from other reports chiefly because we are examining a subpopulation of neurons that were not characterized in previous studies. Alternatively, differences may exist due to distinct bladder layers that were quantified. Some studies physically remove the muscle layers of the bladder from the tissue prior to staining the urothelium and superficial suburothelium (Gabella, 2019; Rahnama’i et al., 2017) while our imaging technique targeted the detrusor muscle and suburothelium layers while omitting the urothelium. One limitation of our study is that in order to quantify nerve density within a standard volume of tissue, we cropped the Z-stacks to include a consistent number of optical slices across all bladders. Based on pilot studies using DAPI to identify changes in the distribution of nuclei between layers we could confirm that our imaging protocol excluded the urothelium, but the amount of suburothelium included in each Z-stack could not be confirmed (data not shown). Thus, it is possible that our image analysis excluded some of the superficial suburothelium from quantification. Although we did not specifically characterize branching patterns, innervation of our bladders appeared consistent with other studies that have found variable patterns ranging from single bundles to more complex bifurcations (Spencer et al., 2018) as well as CGRP fibers with interspaced varicosities (Gabella, 2019).

Collectively, our study provides novel tools to study bladder innervation by non-peptidergic neuronal subtypes without reliance on immunostaining or electrophysiologic properties. For the study of cellular and molecular mechanisms of bladder function, TrkB- and Ret-driven genetic strategies may provide an invaluable tool for the study of mechanosensitive afferent subtypes. These strategies can likewise improve our understanding of the role of mechanoreceptors in bladder pain, bladder overactivity, and other urologic diseases.

## Acknowledgements

We thank Michael Caterina and David Ginty for the founders that were kindly provided to establish the mouse lines utilized in this study.

## Figure Legends

**Supplemental Figure S1. Computational deconvolution allows visualization and quantification of individual nerve terminals**. (A) Diagram illustrating the regions of the bladder that were targeted for imaging and analysis. (B) A raw image of whole-mount immunostained bladder where innervation of the body region appears hazy due to the thickness of the tissue. (Maximum projections, z = 86 with nyquist sampling frequency). (C) After computational clearing of the same image using the large volume computation clearing algorithm on the Leica Thunder 3D imaging system (Thunder regularization for NFH was 4.26e-6 and Thunder strength was 9.8e-1. Both CGRP and mCherry regularization was 4.26e-9 and strength was 9.2e-9. The regularization method used for each channel was Good’s Roughness), innervation patterns are more easily discerned, enabling quantification of fine nerve terminals. (Maximum projection, z = 86 with Nyquist sampling frequency). Scale bar is 100 um. (D) A 3-D reconstruction shows the computationally cleared, full thickness Z-stack from another bladder from a TrkB CreER mouse line. Optical slices from Z-stacks were used to quantify colocalization, in lieu of maximum projections.

**Supplemental Figure S2. Little to no labeling of bladder afferents are seen in Split-CRE mice with labeling characteristic of A-beta low threshold mechanoreceptors**. (A) Because labeling in Split-CRE mice becomes less specific over time in aged mice, formalin-fixed biopsies of the ear skin were evaluated in Split mice to screen for specific labeling of rapidly adapting A-beta low threshold mechanoreceptor skin afferents. Red arrows indicate hair follicles with lanceolate endings, in addition to moderate background labeling of scattered putative keratinocytes. Native tdTomato fluorescence in uncleared, formalin-fixed, whole-mount ear skin. (B) In a mouse with confirmed A-beta mechanoreceptor labeling in the skin, tdTomato labeling was sparse to completely absent in the bladder. (C) Overall the Split mouse line had highly variable tdTomato expression patterns with little to no labeling in bladders from mice with characteristic A-beta mechanoreceptor labeling in the skin (filled circles). In one mouse with abundant non-specific labeling in skin, the bladder had a modest degree of nerve terminal labeling, particularly in the dome (diamonds). N = 3. Scale bars are 100um.

