## Supplemental figures for "Genetic tools that target mechanoreceptors produce reliable labeling of bladder afferents"

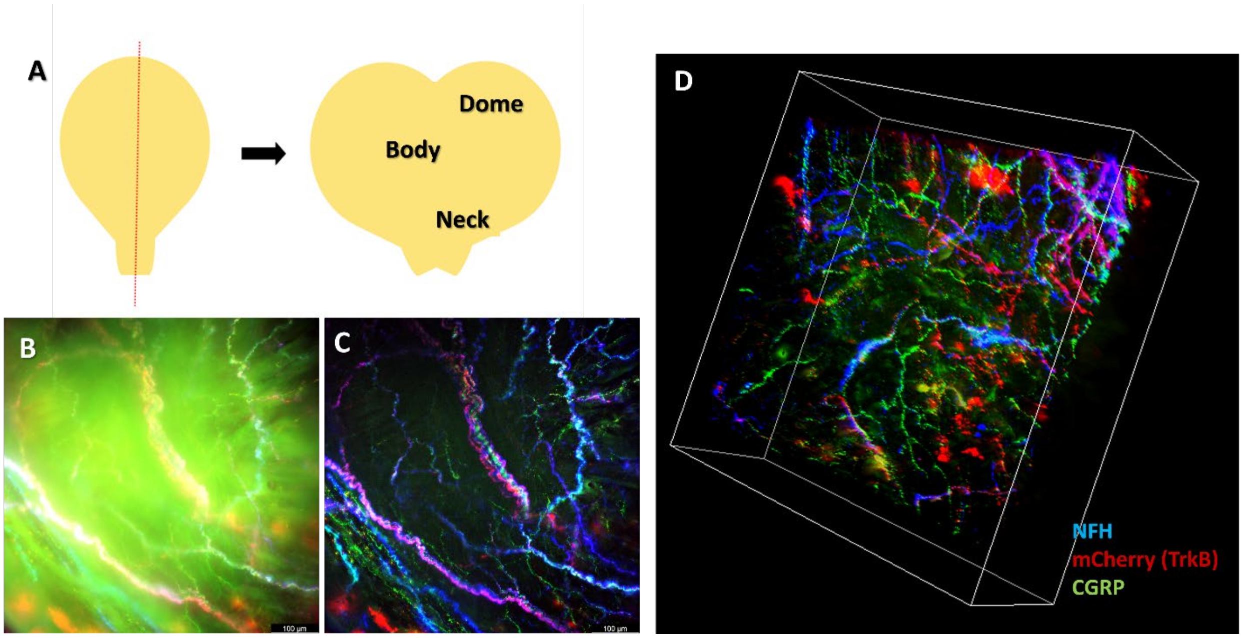


**Supplemental Figure S1. Computational deconvolution allows visualization and quantification of individual nerve terminals.** (A) Diagram illustrating the regions of the bladder that were targeted for imaging and analysis. (B) A raw image of whole-mount immunostained bladder where innervation of the body region appears hazy due to the thickness of the tissue. (Maximum projections, z = 86 with nyquist sampling frequency). (C) After computational clearing of the same image using the large volume computation clearing algorithm on the Leica Thunder 3D imaging system (Thunder regularization for NFH was 4.26e-6 and Thunder strength was 9.8e-1. Both CGRP and mCherry regularization was 4.26e-9 and strength was 9.2e-9. The regularization method used for each channel was Good's Roughness), innervation patterns are more easily discerned, enabling quantification of fine nerve terminals. (Maximum projection, z = 86 with Nyquist sampling frequency). Scale bar is 100 um. (D) A 3-D reconstruction shows the computationally cleared, full thickness Z-stack from another bladder from a TrkB CreER mouse line. Optical slices from Z-stacks were used to quantify colocalization, in lieu of maximum projections.

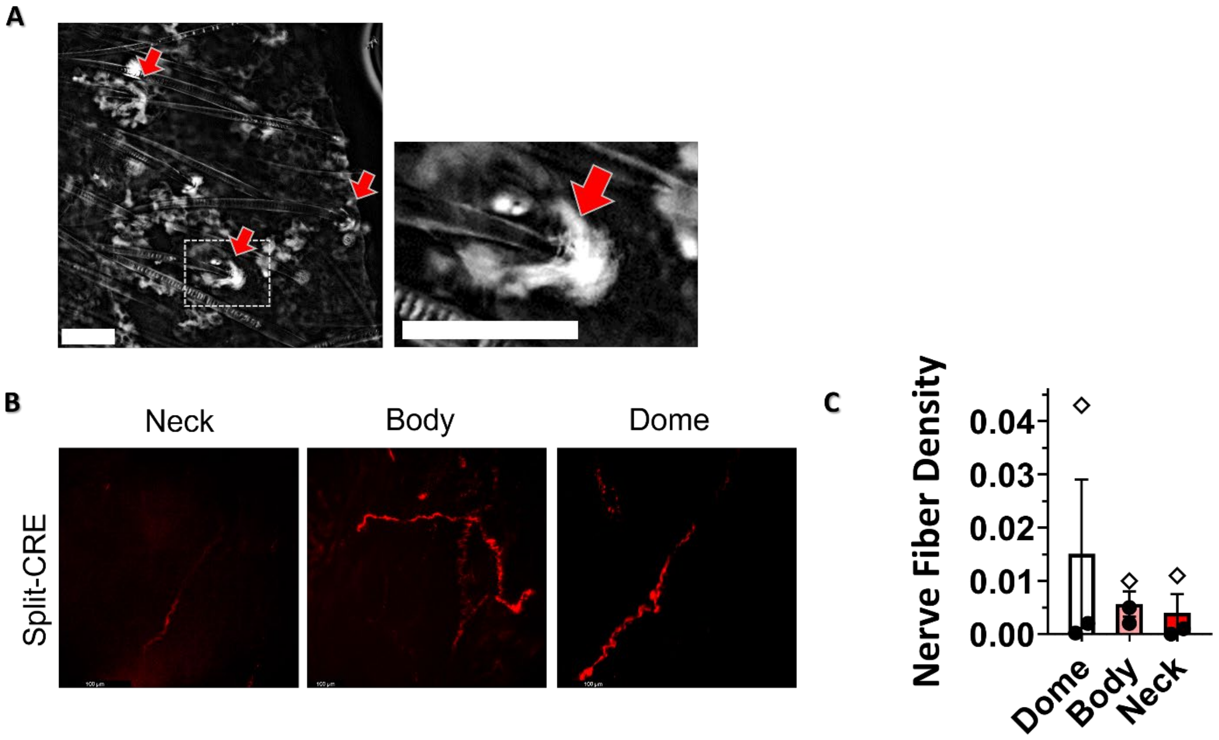


**Supplemental Figure S2. Little to no labeling of bladder afferents are seen in Split-CRE mice with labeling characteristic of A-beta low threshold mechanoreceptors.** (A) Because labeling in Split-CRE mice becomes less specific over time in aged mice, formalin-fixed biopsies of the ear skin were evaluated in Split mice to screen for specific labeling of rapidly adapting A-beta low threshold mechanoreceptor skin afferents. Red arrows indicate hair follicles with lanceolate endings, in addition to moderate background labeling of scattered putative keratinocytes. Native tdTomato fluorescence in uncleared, formalin-fixed, whole-mount ear skin. (B) In a mouse with confirmed A-beta mechanoreceptor labeling in the skin, tdTomato labeling was sparse to completely absent in the bladder. (C) Overall the Split mouse line had highly variable tdTomato expression patterns with little to no labeling in bladders from mice with characteristic A-beta mechanoreceptor labeling in the skin (filled circles). In one mouse with abundant non-specific labeling in skin, the bladder had a modest degree of nerve terminal labeling, particularly in the dome (diamonds). N = 3. Scale bars are 100um.
